# The Histopathological Assesments of Effecting Fe and Zn Heavy Metals in Cerebrum and Cerebellum Tissues of Rats

**DOI:** 10.1101/2022.06.21.496962

**Authors:** N. Şenol, M. Şahin

## Abstract

Heavy metals left intensively to the environment cause different healthy problems on people by entering the food chain heavy metals. Our aim is to provide determinig effect of in tissues especially in central nervous system as a result of consuming cause foods included heavy metals and water. Five groups were constituded by using 35 adult male Wistar-Albino sexual rats. Hematoxylin-eosin (H&E) staining was applied to determine the histological sides of the damages of heavy metals in cerebrum and cerebellum tissues and effects of given juglone (5-hydroxy-1,4-naphthoquinone) for reducing these damages. Besides, immunohistochemical TUNNEL method was applied to determine DNA damages in cell. Density of damage in cerebrum of Fe and Fe+Juglone groups were higher than in the control group. Apoptotic cells number was decreased in the group consisting of juglone. There was little damage and less apoptotic cells in treated Zn group. In the cerebellum similar cerebrum results were obtained. There were much degeneration Fe and Fe+Juglone groups cerebellum. Apoptotic cells were increased Fe groups. Fe/600 ppm. doses heavy metals have created a toxic effect on the cerebrum and cerebellum tissues.

## Introduction

Most of the animal and neurotoxic effects of heavy metals and peripheral nervous system indicated major involvement of blood vessels. However, little is known of its mechanism of neurotoxic action as well as the comparative evaluation of the morphological lesions after identical exposure in developing and adult brain (Murthy et al. 1987). Iron is an essential mineral in humans and plays a crucial role in vital biochemical activities, such as oxygen sensing and transport, electron transfer and catalysis (Kozan et al. 2008).

Heavy metals are naturally occurring metal that is present in food, soil and water. It is relased in to the enviroment from both natural and man-made sources. Chronic exposure to heavy metal-contaminated water and food causes cancer of skin, liver, lung and bladder. Heavy metal is also said to exert its toxicity through oxidative stress by generating reactive oxygen species. Free radicals have been detected in some cell treated with heavy metal. Treament with heavy metal has been shown to induce of hydroxyl radical formation in brain (Zalups and Ahmad 2003; Bridges and Zalups 2005; Bashir et al. 2006).

Heavy metals are the most important source of anorganic pollution in freshwater. Heavy metals are transported by rock pieces transported with erosion, dust transported by wing, volcanic activities, forest fires and plant cover to the water. Chemical pollutants involved to aquatic environment via atmosphere. Because these elements which are in atmosphere get to the water with the rain and wind and has an impact on the aquatic systems (Lee et al. 2007; Sönmez and Akkuş 2009).

Heavy metals left intensively to the environment cause different healthy problems on people by entering the food chain heavy metals. The food chain far reaching people through heavy metals in humans also causes a number of diseases and even death. Our aim is to provide determinig effect of in tissues especially in central nervous system as a result of consuming cause foods included heavy metals and water. Because of this study, the male rat specific ppm-level heavy metals (Fe and Zn) effects on brain tissue damages created recursively, and to reduce the effects of antioxidants (Juglone (5-hydroxy-1,4-naphthoquinone) were determined as the histological and immunohistochemical applications.

## Materials and Methods

The damage of heavy metals, iron and zinc formed in the rats brain tissue and juglone (5-hydroxy-1,4-naphthoquinone) antioxidant activities in preventing these damages were explored with histological and immunohistochemical methods in this study. Five groups were constituded by using 35 adult male Wistar-Albino sexual rats. First group was control group (1 ml. water), second was given iron (0.3 ml. stock solution from Fe/600 ppm. + 0.7 ml.water), third was given Zinc (0.2 ml. stock solution of Zn/ 400 ppm+ 0.8 ml. water), fourth was given Fe (0.3 ml. stock solution from Fe/600 ppm. + 0.7 ml.water) +Antioxidant Juglone (5-hydroxy-1,4-naphthoquinone), fifth was given Zn (0.2 ml. stock solution of Zn/ 400 ppm) + Antioxidant Juglone (5-hydroxy-1,4-naphthoquinone) which was given to the method of rat Gavage. Hematoxylin-eosin (H&E) staining was applied to determine the histological sides of the damages of heavy metals in cerebrum and cerebellum tissues and effects of given juglone (5-hydroxy-1,4-naphthoquinone) for reducing these damages. Tissues were observed with light microscopic (Leica DM 500) examinations. Besides, immunohistochemical TUNNEL method was applied to determine DNA damages in cell (Bancroft et al. 2008).

## Results

It was observed that the number of ischemic neurons and vascular dilatation were more density in Fe group. Density of damage in brain tissue of Fe group was higher than in the control group (Figure1-4) but in the other groups were no significant difference with control group (p□0.05). Vacuolization in neurons were identificated at Fe group. Number of apoptotic cells were observed in each group. Density of damage in cerebrum of Fe and Fe+Juglone groups were higher than in the control group. Apoptotic cells were increased Fe groups (Figure 4). Apoptotic cells number was decreased in the group consisting of juglone. There was little damage and less apoptotic cells in treated Zn group.

**Fig. 1:**
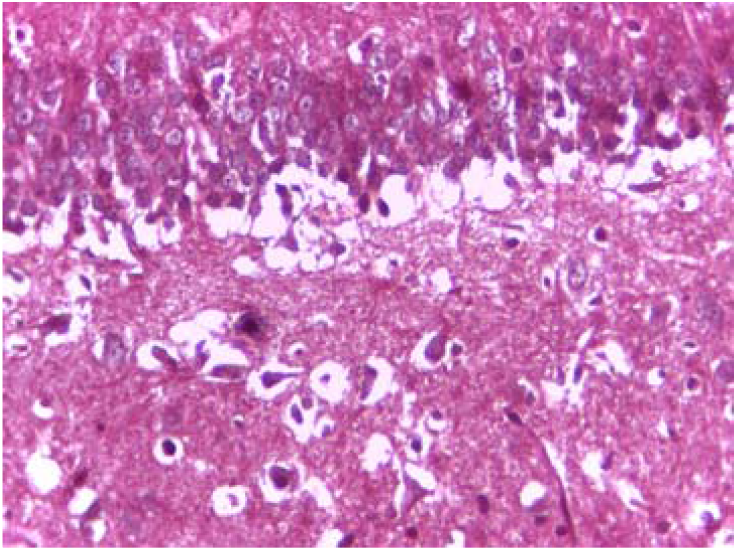
Fe Group, Masson trichrome, X 40

**Fig. 2:**
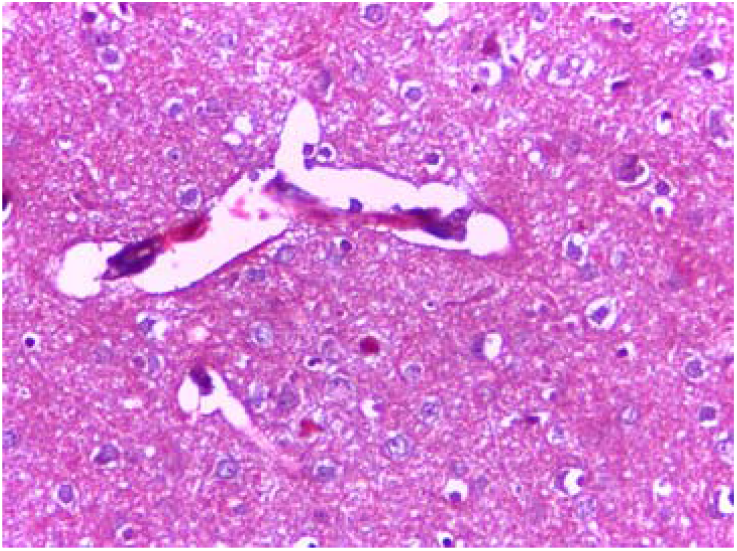
Fe Group, Masson trichrome, X 40

**Fig. 3:**
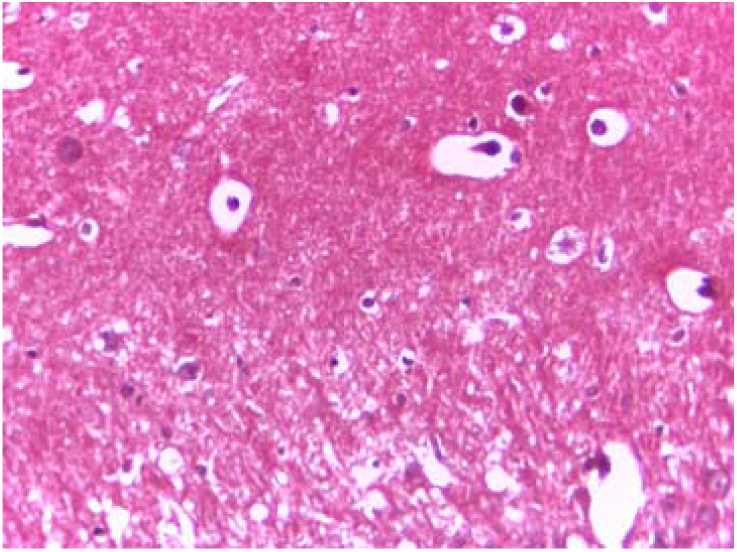
Fe Group, Masson trichrome, X 40

**Fig. 4:**
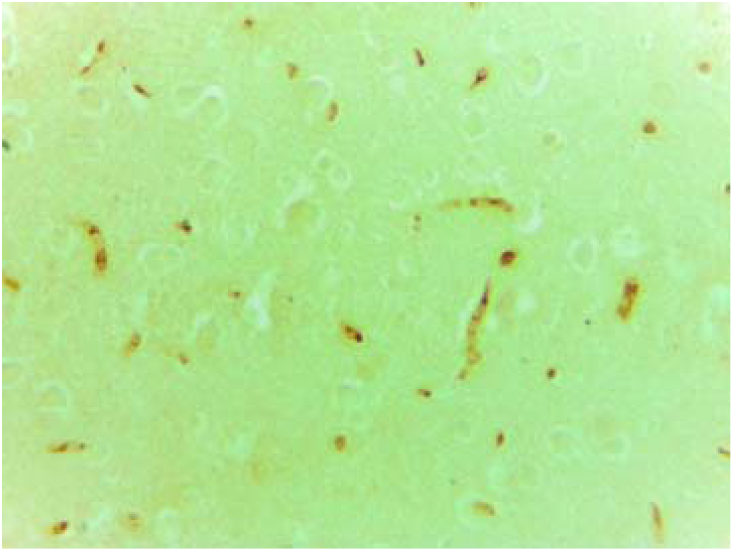
Fe Group, TUNEL, X 40

Number of apoptotic cells and purkinje cells were observed in each group. Density of damage in cerebellum of Fe and Fe+Juglone groups were higher than in the control group but in the other groups were no significant difference with control group (p□0.05) (Figure 5-10). Degeneration and reduction in the number of purkinje cells were determined in Fe and Fe+Juglone groups. Apoptotic cells were increased Fe groups.Degeneration in Fe+Juglone group was decreased according to the Fe group because of protective effect of the juglone.

**Fig. 5:**
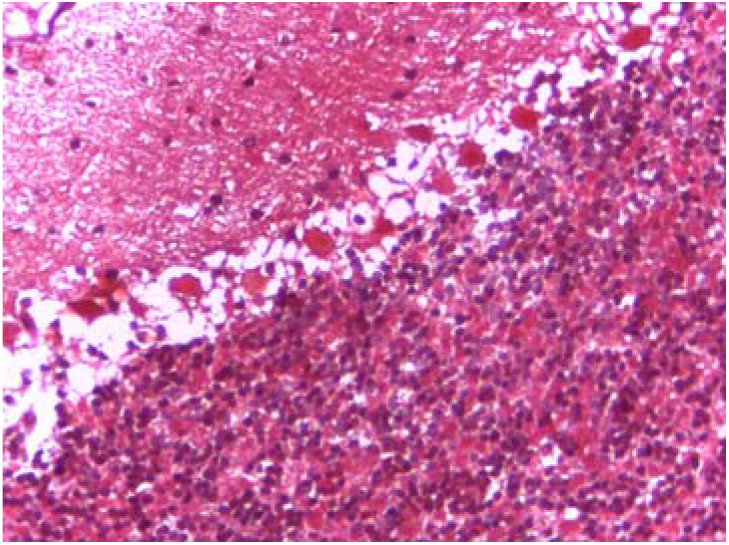
Fe Group, Masson trichrome, X 40

**Fig. 6:**
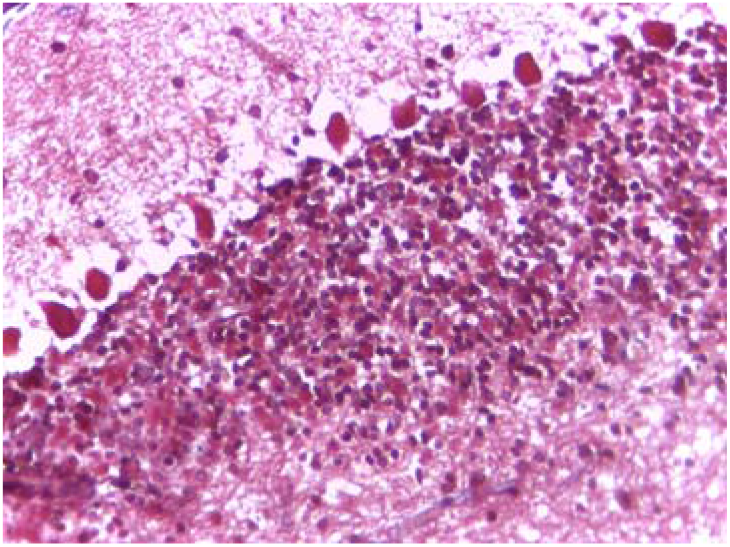
Fe+Juglone Group, Masson trichrome, X 40

**Fig. 7:**
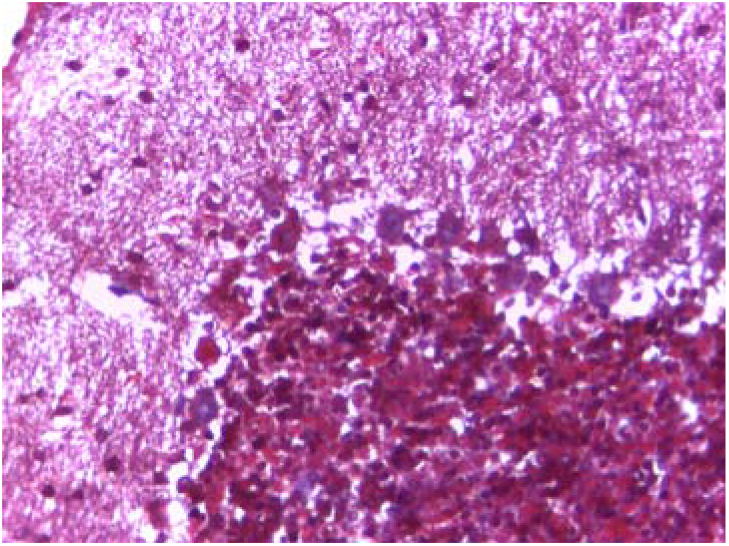
Zn Group, Masson trichrome, X 40

**Fig. 8:**
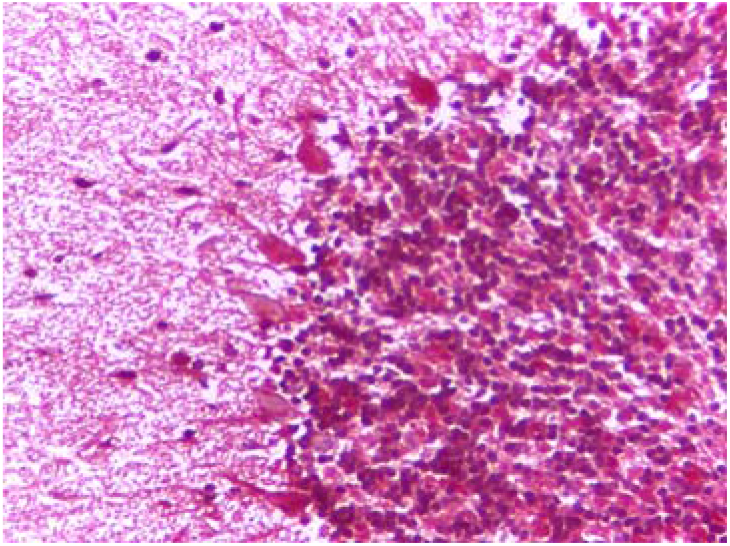
Zn+Juglone Group, Masson trichrome, X 40

**Fig. 9:**
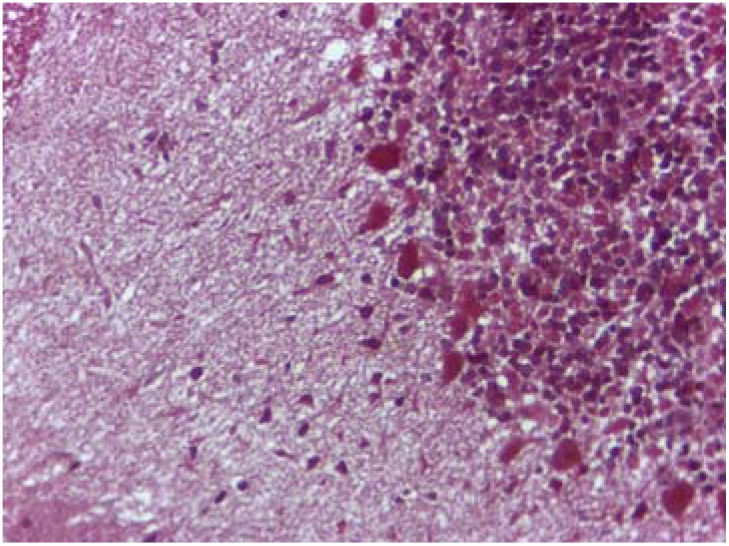
Control Group, Masson trichrome, X 40

**Fig. 10:**
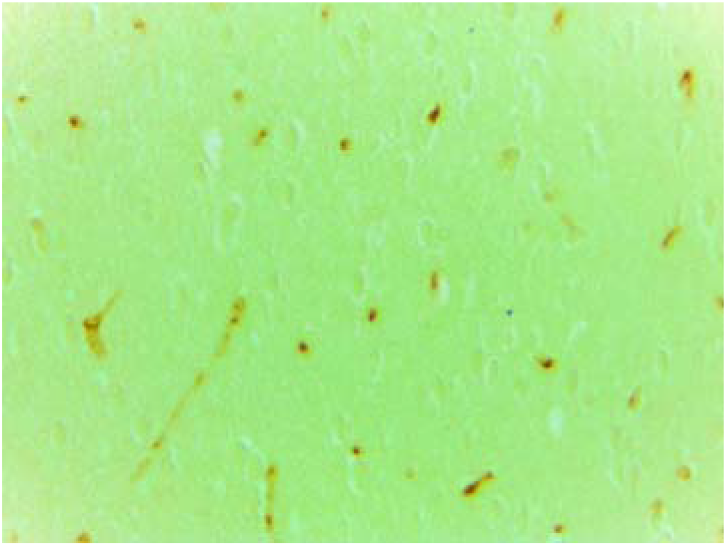
Fe Group, TUNEL, X 40

## Discussion

Various heavy metals such as cadmium, copper, mercury, iron and zinc are well documented to stimulate, tissue metallothionein. Metallothionein (MT) is found in various animal tissues, including the brain, and functions as a scavenger of heavy metals as well as a survey for essential metals (Yasutake et al. 2004).

In spite of exposure to mercury vapor for 3 months, no pathological change was determined in the brains of either MT-null or wild type mice, suggesting a low mercury toxicity of the present dose schedule. The presence of mercury could be detected histochemically in the brains of grains were observed in the cytoplasm of nerve cells and occasional glial cells throughout the brain of both strains, though non-exposed control tissues indicated no mercury grains. It was not observed differences in the distribution and intensity of mercury deposits between the brains of MT-null and wild type mice exposed to mercury (Yasutake et al. 2004). It was reported that mercury deposition at cerebrum and cerebellum cells were observed in rats which exposed to mercury 10 weeks (Warfvinge 1995).

Nehru et al. (1997) were investigated when the weight of the cerebrum and cerebellum was taken separately, a significant rise in the cerebellar weight was seen in the Pb-treated group, and it resulted in a important increase in its rate to brain weight. Following combined application of Pb and Se a substantial improvement in the cerebrum weight and brain weight was declared. Histologically, the transverse section of cortex of group I (control) animals indicated a well-organized cortical layer. The cells were uniform and showed no vacuoles. In the Pb-treated animals, however, the layers were almost absent, and the neurons were diffuse in the cross-section. There were spaces and autophagic vacuoles with debris seen in many places. Pb and Se treated animals, the autophagic vacuoles were present. The pyramidal cells were reduces in size. In this study, it was observed that the number of ischemic neurons and vascular dilatation were more density in Fe group. Density of damage in brain tissue of Fe group was higher than in the control group but in the other groups were no significant difference with control group.

In the central nervous system signs of silver were identicated after higher doses of silver lactate. The sediment was confined to lysosomes of motor neurons in the pontine nuclei and in the spinal cord, but glial cells were also found to contain silver. Neuropathic changes were seen in the myelin sheaths of cauda equina axons (Danscher 1991).

Histochemical changes in several neurons in the central nervous system and spinal ganglia could be detected after about 14 days of exposure to mercury chloride. By investigation light microscopy, the reaction products were seen as black grains in the neuronal somata. In many cases the grains were localized to special regioni juxtanuclearly, but more even distributions were also found (Danscher and Schroder 1979).

Total number of Purkinje cells in iron and iron+nicardipine groups were significantly lower than control animals (p<0.005). However, when the iron and iron+nicardipine groups were compared, purkinje cell loss was higher in the iron group (p<0.05) (Kozan et al. 2008). Similar results were obtained in this study.

The administration of cadmium (100 ppm) in drinking water to growing rats from 21 days of life for 120 days consistantly induced lesions in the cerebellar cortex. Purkinje cells at places were found to be disintegrated with pyknotic nuclei, eosinophilic cytoplasm and resolution of cellular membranes. The blood vessels, however, appeared normal. The leptomeninges and deep cerebellar nuclei were splitted up (Murthy et al. 1987).

Similar results were observed cerebellum tissues. Density of damage in cerebellum of Fe and Fe+Juglone groups were higher than in the control group but in the other groups were no significant difference with control group (p□0.05). Degeneration and reduction in the number of purkinje cells were determined in Fe and Fe+Juglone groups.

Arsenic dose-dependent histopathological changes observed in brain. The section from brain show more frequent nuclear pyknosis. A significant increase in caspase-3 activity was determined at 10.5 and 12.6 mg/kg sodium arsenite in brain (Bashir et al. 2006). In this study, apoptotic cells number was higher Fe groups than other groups.

Fe/600 ppm.doses heavy metals have created a toxic effect on the cerebrum and cerebellum tissues. The damage associated with the Zn and Zn+Juglone was determined to be less significant than the damage by Fe and Fe+Juglone groups. Degeneration in cerebrum and cerebellum tissues were decreased with antioxidant effects of juglone.

## Authors’ contribution

NŞ and MŞ conceived and planned the study. NŞ drafted and revised the manuscript. NŞ collected the data. NŞ and MŞ analyzed the data. All authors approved the final version of the manuscript for submission.

## Data availability statement

All the research data related to this manuscript will be available upon reasonable request to the corresponding authors.

## Declaration of conflicting interests

The author(s) declared no potential conflicts of interest with respect to the research, authorship, and/or publication of this article.

## Ethical approval

The ethical approval of the study was obtained from the Süleyman Demirel University Animal Experiments Local Ethics Committee Chairman Turkey (21438139-338).

## Funding

The author(s) disclosed receipt of the following financial support for the research, authorship, and/or publication of this article: This research was funded by Scientific and Technological Research Council of Turkey (Project number: 214O699).

## Supplemental material

Supplemental material for this article is available online.

## Notes

### Competing Interest Statement

The authors have declared no competing interest.

